# The Establishment and Potential Spread of *Osmia cornuta* (Hymenoptera: Megachilidae) in North America

**DOI:** 10.1101/2024.06.18.599626

**Authors:** Michael P Getz, Lincoln R Best, Andony P Melathopoulos, Timothy L Warren

## Abstract

Mason bees, subgenus *Osmia* Panzer (Hymenoptera: Megachilidae), are economically and ecologically significant pollinators. In eastern North America, the rapid spread of two non-native species from Asia, *O. cornifrons* Radoszkowski and *O. taurus* Smith, has coincided with declines in native *Osmia* populations, raising concern about the effects of further exotic invasions. Here we investigate the recent establishment in British Columbia, Canada of the European orchard bee, *O. cornuta* Latreille, previously thought to be limited to Europe and its periphery. We document *O. cornuta* records ranging over 170 km, including sightings of live adults and the discovery of a multigenerational nest with hundreds of cocoons. We tested whether these *O. cornuta* cocoons could be discriminated from other *Osmia* species by training a machine learning classifier on features extracted from images. The best performing model could not reliably discriminate cocoons by species, raising the possibility *O. cornuta* could be inadvertently intermingled in future commercial shipments. Spatially isolated records of *O. cornifrons* and *O. taurus* further suggest ongoing anthropogenic dispersal of these species. To determine environmentally suitable regions for *O. cornuta* to spread in North America, we estimated its climate niche using native-range occurrence data. This analysis indicated broad regions of the Pacific Northwest and eastern North America contain potentially suitable habitat. Together, our findings document the establishment of *O. cornuta* in North America and the potential for it to spread broadly. Our study demonstrates the utility of accessible biodiversity data archives and public observation programs in tracking biological invasions and highlights the need for future monitoring of exotic *Osmia*.

## Introduction

The mason bees, subgenus *Osmia* (*Osmia*) Panzer (Hymenoptera: Megachilidae), hereafter *Osmia* s.s., are economically and ecologically significant pollinators that are widely distributed throughout the Holarctic region. Of the 29 bees in the subgenus, several have become essential managed pollinators for agricultural systems in their native ranges, including *O. lignaria* Say in North America, *O. cornifrons* Radoszkowski in northern Asia, and *O. cornuta* Latreille in Europe (Branstetter et al. 2021). Mason bees are ideal for husbandry and agricultural use, as they can be more effective pollinators of rosaceous trees than *Apis mellifera* L. and *Bombus terrestris* L. (Bosch and Blas 1994, Jaumejoan et al. 2023, Vicens and Bosch 2000), they readily build nests in provided artificial cavities, and are easily transported in their cocoon stage (Sedivy and Dorn 2014, Torchio et al. 1987). In recent decades, two non-native *Osmia* s.s., *O. cornifrons* and *O. taurus* Smith, have spread widely in eastern North America. *O. cornifrons* was first established in Maryland as a result of a prolonged, intentional introduction effort, involving multiple shipments of thousands of bees from Japan as early as 1965 (Batra 1998, Rust 1974). In contrast, the closely related *O. taurus*, also native to Japan, was introduced unintentionally to the eastern US; its first documented records in North America were in Maryland and West Virginia in 2002 (Droege and Maffei 2023). Both species have broadly increased their range in the last 20 years. Their shared range now extends from the Carolinas to southern Ontario and west into several midwestern states (Gutierrez et al. 2023, MacIvor et al. 2022). *Osmia taurus* observations reach as far south as Georgia and Florida, and *O. cornifrons* records extend west to Colorado, Utah, and Oregon (Gutierrez et al. 2023).

There is growing concern about the extent to which exotic *Osmia* species could threaten native ecosystems. In a multidecade monitoring study in the mid-Atlantic US, annual increases in *O. cornifrons* and *O. taurus* abundance coincided with significant declines in all six native *Osmia* species tracked, including *O. lignaria* (LeCroy et al. 2020). One possible cause for these apparent population declines is transmission of fungal pathogens from non-native to native species (Crowl et al. 2008, Prenter et al. 2004). *O. cornifrons* populations in the US have been shown to harbor fungal pathogens from Japan (Hedtke et al. 2015), whereas natives *O. lignaria* and *O. georgica* have been found carrying chalkbrood-causing fungi (*A. naganensis* and *A. fusiformis*) originally characterized in Japan (LeCroy et al. 2023). In addition to transmitting disease, exotic bees can compete with native species for food and nesting resources and alter plant communities through selective pollination (Goulson 2003, Russo et al. 2021, Stout and Morales 2009, Thomson 2004).

*Osmia* s.s. have several attributes that contribute to their success as invasive species. As cavity nesters, they can disperse globally by building nests in materials used for international shipping (Goulson 2003). *Osmia* s.s. are generally polylectic, which supports flexible adaptation to new environments (Haider et al. 2014), although some species have known floral preferences (e.g. *O. ribifloris* for *Arctostaphylos*). Exotic *Osmia* can benefit from commercial beekeeping operations, which provide housing, food, and further opportunity for anthropogenic dispersal. In particular, it has been hypothesized that the shipment of cocoons – either by large commercial operators or by small-scale hobbyists – can accelerate the inadvertent dispersal of *Osmia* s.s. (MacIvor et al. 2022). Although some states regulate the import of non-native insects, there are currently no federal limits on domestic shipments of *Osmia* s.s within the continental US or Canada (USDA 2018).

Despite the high invasive capacity of *Osmia s*.*s*., many intentional introductions have failed, suggesting that habitat suitability critically limits the viability of establishment. For instance, prior to the successful establishment of *O. cornifrons* to Maryland in 1977, an attempted introduction to Utah failed in 1965. When researchers released thousands of European orchard bees (*O. cornuta*) in Logan, Utah in 1978, they assumed the species would establish permanently (Torchio and Asensio 1985). Several *O. cornuta* nests were found the following year, but there were no records of the species thereafter (Torchio and Asensio 1985). The same group then attempted to introduce *O. cornuta* to four almond orchards in Dixon, California in 1985; again, there was no lasting establishment (Torchio and Asensio 1985, Torchio et al. 1987). These failed introductions may have resulted from inhospitable climate conditions. As univoltine bees, *Osmia* s.s. are highly sensitive to the quantity and timing of rainfall, which affects spring emergence and foraging success, and to seasonal temperature variation, which influences developmental rate and the viability of overwintering (Kemp and Bosch 2005, Pitts-Singer et al. 2014, Radmacher and Strohm 2011, Westreich et al. 2023, White et al. 2009). Consistent with this environmental sensitivity, *O. lignaria* survived at a higher rate when introduced to locations with climates that closely matched their original range (Kemp and Bosch 2005). Therefore, as for many invasive taxa, determining the climatic niche of exotic *Osmia* species is an essential part of predicting their spread (De Marco and Nóbrega 2018, Polidori and Sánchez-Fernández 2020, Silva et al. 2014). To model habitat niche at a broad scale, many studies have used the abundant presence-only data generated by community science monitoring (Di Febbraro et al. 2023, Encarnação et al. 2021, Tiago et al. 2017).

The successful spread of *O. cornifrons* and *O. taurus* raises the possibility that other exotic *Osmia* could similarly establish in North America with potential negative consequences for native bees. This study investigates the recent establishment of *O. cornuta* – thought to be limited to Europe and its periphery – in southern British Columbia, Canada. We document numerous sightings of *O. cornuta* adults as well as a multigenerational nest with hundreds of cocoons. We evaluate the visual features of these cocoons to assess whether *O. cornuta* could go undetected in shipments of other *Osmia* s.s. cocoons. To predict *O. cornuta*’s range expansion dynamics, we examine the historical spread of other non-native *Osmia* s.s. and estimate its native climate niche.

## Methods

### Sample collection and occurrence records

In April 2023, Gordon Cyr, the owner and operator of a mason bee farm in Black Creek, Vancouver Island, harvested a cache of cocoons from a prefabricated mason bee house after observing distinctly red adult mason bees nearby. Suspecting they were *O. cornuta*, Cyr contacted the Native Bee Society of British Columbia (NBSBC), which arranged for the transport of 30 cocoons to Oregon State University (OSU) – suspended in ethyl alcohol – for identification and dissection. In January 2024, Cyr harvested 80 cocoons from the same nesting site, which he again sent to OSU. We obtained *O. lignaria* and *O. cornifrons* cocoons from Willamette Valley beekeepers as part of an Oregon mason bee health project. All cocoons were coarsely cleaned of nesting materials, with occasional frass and mud cap fragments remaining.

We accessed four iNaturalist records of adult *O. cornuta* from British Columbia, Canada: a March 29, 2023 record from North Vancouver, BC (153554196) contributed by Jasna Guy, a Master Melittologist volunteer for the NBSBC; two concurrent observations (204882854, 204883055) of a male and a female *O. cornuta* on March 30, 2024 contributed by Jeremy Gatten, Wildlife Biologist, LGL Limited; and an April 15, 2024 record of a female *O. cornuta* (207021500) contributed by Bonnie Zand, a director and taxonomist for the NBSBC. We visually verified the identifications made for all photos.

### Cocoon and specimen analysis

We imaged 317 cocoons from three species, *O. lignaria* (n = 130), *O. cornifrons* (n = 81), and *O. cornuta* (n = 106) using a Raspberry Pi 4 with a Pi Cam HQ 12MP module, 6mm lens (Adafruit) at a constant height (20 mm), and an LED ring light. After imaging we dissected the cocoons and identified specimens to species and sex (Amiet et al. 2004), and measured intertegular distances using digital calipers to 0.01 mm accuracy under a dissecting scope (Leica). We implemented the analysis of cocoon images in Python. We corrected for lens distortion using the OpenCV Camera Calibration procedure (Bradski 2000). We segmented cocoons in images by first training a YOLO object detection model (yolov8s) and used the resulting bounding boxes as inputs for Segment-Anything (Meta; Kirillov et al. 2023, Jocher et al. 2023). We calculated cocoon length and width with OpenCV by fitting a rectangle to the resulting cocoon mask. Area (cm^2^) and average color values on a scale of 0 to 255 were extracted from the cocoon mask. To estimate discriminability among cocoons from visual characteristics, we used the scikit-learn package to train a support vector machine (SVM) model on the extracted features (length, width, area, and red, green, and blue color values; Pedregosa et al. 2011). We used a 10-fold cross-validated grid search to tune SVM parameters: gamma (influence range for each training example), C (complexity of decision boundary), and kernel type (linear, radial basis function, or polynomial). Using the best parameters from the grid search (C = 100, gamma = 0.1, kernel = radial basis function) we performed 1000 iterations of 10-fold cross-validation. We tested two conditions using a one-vs-rest format: whether *O. cornuta* could be reliably classified to prevent its spread or detect its arrival in a new area, and whether *O. lignaria* cocoons could be isolated from non-natives. We evaluated the performance of the SVM by averaging precision and recall across all iterations. For a particular class (e.g. *O. lignaria* or *O. cornuta*), precision is the ratio of true positives to total positive determinations. Recall is the proportion of actual positives correctly classified (i.e. true positives divided by true positives and false negatives).

### Distribution mapping

We generated historical species distribution maps in Python using the Global Biodiversity Information Facility (GBIF) database, a repository for peer-confirmed research data and community scientist submissions. A “Research Grade” observation requires a date, georeferenced coordinates, an associated photo or sound, at least two confirmatory identifications to species level or lower, and at least two-thirds of identifications must agree. For *O. lignaria, O. cornifrons*, and *O. taurus*, we mapped the locations of all North American GBIF records from 1960-2023 (between 23 to 60 deg latitude and -126 to -64 deg longitude; GBIF 2024). We projected a grid of equally sized bins (22,500 km^2^) over this area, with the projection centered at -95 deg longitude. For each species on a yearly basis, we tracked the total quantity of observations made and the number of unique spatial bins with at least one record. We further tracked the cumulative number of bins that contained a species record for the first time that year, to determine whether species were being observed in new areas.

### Assessing habitat similarity

We assessed the suitability of North American habitat for *O. cornuta* using R, first by estimating the environmental niche of *O. cornuta* in its native range (i.e. the range of climate conditions in which it was observed). At each of the 32,413 *O. cornuta* occurrence locations (1960-2023; excluding recent North American observations), we extracted values for the 19 bioclimatic variables and elevation from the WorldClim database (Fick and Hijmans 2017, GBIF 2024). These 19 WorldClim variables, derived from temperature or precipitation measures, capture general conditions as well as seasonal trends (Kriticos et al. 2012, Fick and Hijmans 2017). To reduce the effects of uneven sampling density, we filtered the full set of occurrence data to include at most one record per pixel at the resolution of the WorldClim dataset (30 arc seconds, ∼1 km^2^), resulting in 6,494 points. All data were z-score normalized, so that each feature distribution had zero mean and matched standard deviation. We then applied principal components analysis (PCA) to the native-range data to transform the 20 environmental variables, which are highly collinear, to four orthogonal components used for subsequent analysis (De Marco and Nóbrega 2018). These first four components, chosen on the basis of scree plot inspection, accounted for 85.2% of variance. The most positive loadings of the first principal component included maximum temperature of the warmest month and mean temperature of the warmest quarter, whereas the most negative loadings included annual precipitation and precipitation of the driest month. Variables with the most positive loadings on the second axis included temperature seasonality and annual range, whereas minimum temperature of the coldest month and mean temperature of the coldest quarter had the most negative loadings. We estimated *O. cornuta*’s environmental niche by calculating the Mahalanobis distance (*D*^2^) in this four-dimensional PCA space from each of the 6,494 locations to the centroid of these points.

We then extracted the same 20 environmental variables across North America at the full resolution of the WorldClim dataset (∼1 km^2^; from -130 to -63 deg longitude and 25 and 54 deg latitude). We mapped these data (normalized using the native-range means and standard deviations) to the native-range PCA feature space. Then, for each point in North America, we calculated *D*^2^ to the native-range centroid. Smaller *D*^2^ values implied higher similarity to the native climate niche (Farber and Kadmon 2003, Etherington et al. 2009).

## Results

Prior to 2023, the range of the European orchard bee (*O. cornuta*) was limited to continental Europe and its immediate periphery, including the Mediterranean islands, North Africa, and far western Asia. From 1960-2023, 32,416 *O. cornuta* records were logged to the Global Biodiversity Information Facility (GBIF) database (Fig. 1A). The highest density of observations was in central Europe (e.g. Belgium, Holland, and Germany); the spatial range of records extended from Lagoudera, Cyprus to Oslo, Norway (from south to north), and from the Caspian coast in Iran to Serra da Estrela Nature Park, Portugal (from east to west).

**Figure 1:**
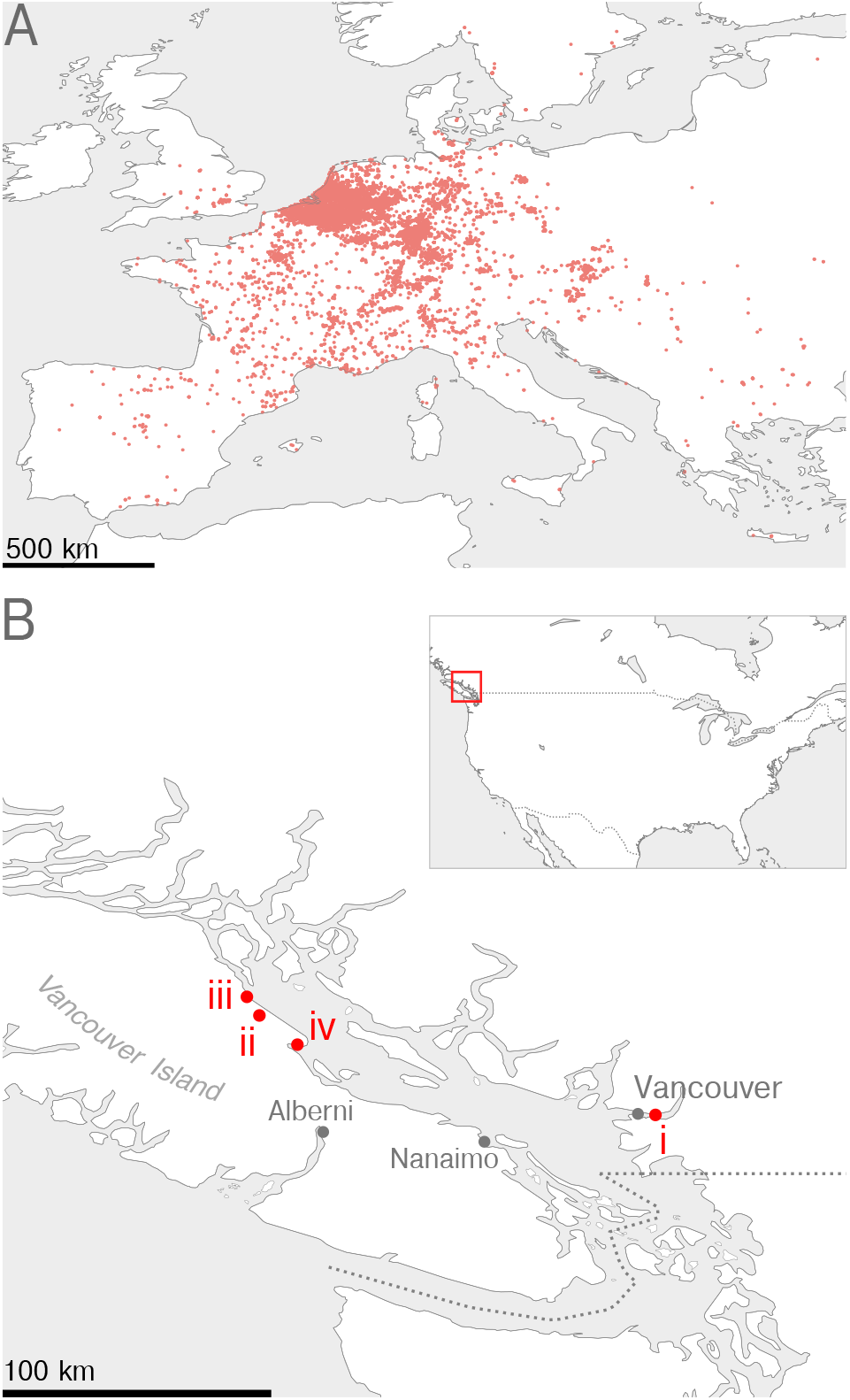
Occurrence records of the European orchard bee, *O. cornuta*, in Europe and North America. A) Locations of 32,295 GBIF occurrence records (red circles) for *O. cornuta* from 1960-2023. Not shown are 118 records to the east between 28.0 to 51.1 degrees longitude. B) Red circles denote four distinct records of *O. cornuta* in British Columbia, Canada in 2023 and 2024: i) an adult male observed in spring 2023 in North Vancouver, BC, ii) a nest with cocoons found in 2023 and 2024 in Black Creek, Vancouver Island, iii) a male and female *O. cornuta* sharing a nest with *O. lignaria*, iv) an adult female *O. cornuta*.

In 2023 and 2024, there were multiple observations of *O. cornuta* in British Columbia, the westernmost province of Canada. In March 2023, a volunteer from the NBSBC encountered an isolated male *O. cornuta* at a residential park in North Vancouver, BC. The image record, uploaded to iNaturalist, was the first valid GBIF entry of *O. cornuta* in North America. In April 2023, in Black Creek, Vancouver Island (Fig 1B; 160 km from North Vancouver), a beekeeper harvested a set of cocoons from a nest where he had observed putative *O. cornuta* adults. We dissected 30 cocoons to determine species and sex; all were *O. cornuta* (11 male, 19 female). Sex was determined by the 13 antennal segments and 7 exposed tergites of males, and 12 antennal segments and 6 exposed tergites of females. Males were identified to species by their tergite 7 with wide, rounded lateral lobes, tergite 6 lacking posterolateral teeth or angles, black pubescence of mesepisternum, and rufescent dorsum of the metasoma (Fig. 2A-C). Females were identified to species by their red metasomal scopa, concave clypeus with laterally protruding horns, black vestiture of head and thorax, and red metasomal tergites (Fig. 2D-F). Intertegular distances of adults (male, mean 3.3 +/- 0.42 mm; female, mean 3.8 +/-0.22 mm) matched prior European estimates and were higher than the respective sexes of *O. lignaria* and *O. cornifrons* samples (Fig. 2G; Bosch and Vicens 2002). We further examined a second batch of 80 cocoons from the same location in 2024. 77/80 were *O. cornuta* (57 male, 20 female).

**Figure 2:**
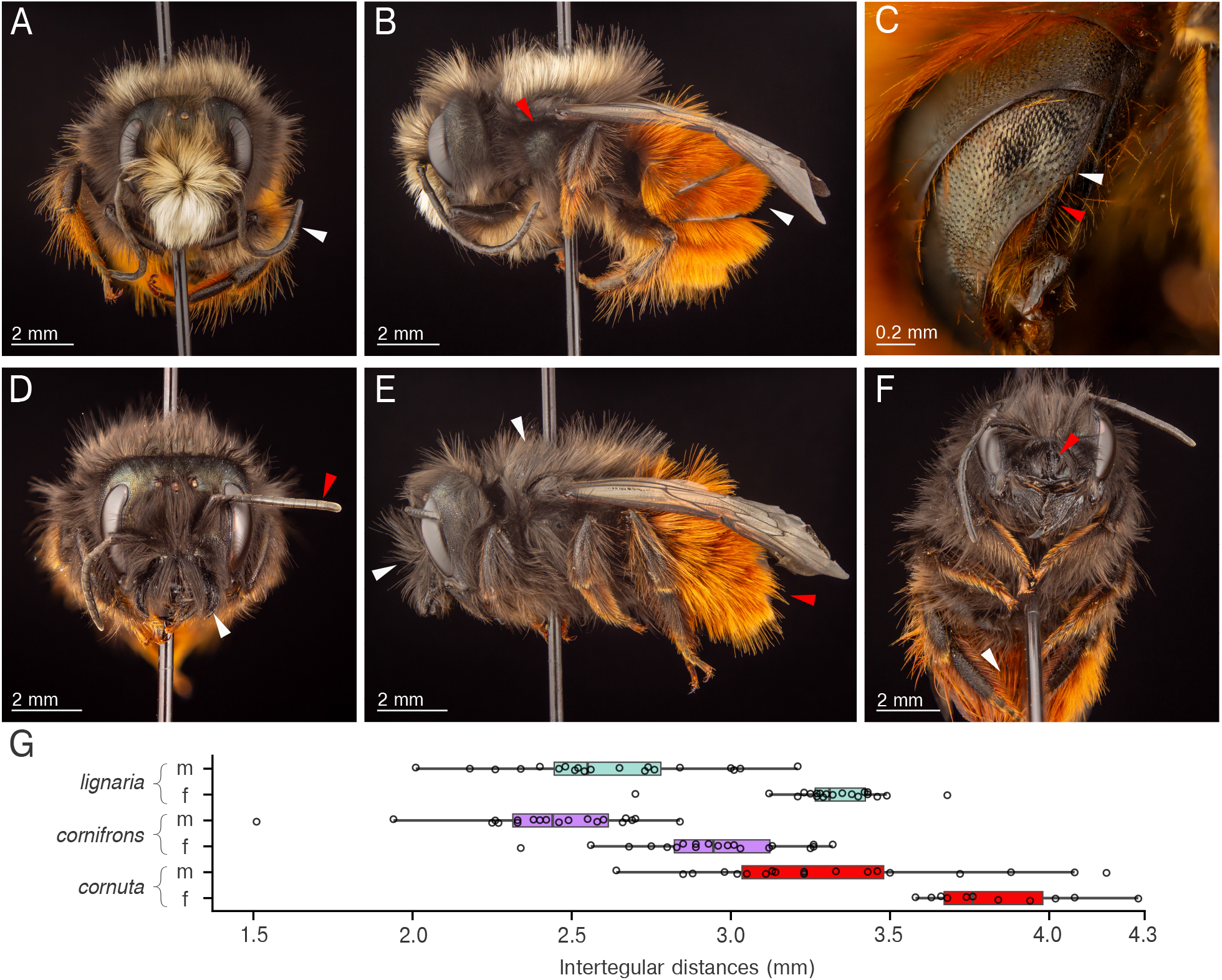
Physical characteristics of Black Creek *O. cornuta* specimens. A) Male face with 11 flagellar segments (white arrow). B) Male lateral habitus with black mesepisternum (red arrow) and rufescent metasoma (white arrow). C) Male tergite 6 & tergite 7 detail (hairs shaved for clarity). Tergite 7 is wide and rounded laterally and lacks teeth (red arrow), whereas tergite 6 lacks angles and teeth (white arrow). D) Female face with 10 flagellar segments (red arrow) and lateral horns (white arrow). E) Female lateral habitus with rufescent metasoma (red arrow) and black head and mesosoma (white arrows). F) Female ventral habitus with concave clypeus (red arrow) and red scopal hairs (white arrow). G) Box and whisker plot for the intertegular distances of *O. cornuta, O. lignaria*, and *O. cornifrons*. Boxes mark the interquartile range (IQR) from Q1 to Q3 and median. The whiskers mark the minimum and maximum values within 1.5 times the IQR. Open circles are measured values; circles outside the whiskers are considered outliers. Image credit: Mark Gorman

One cocoon was *O. lignaria* and two contained underdeveloped larvae. In spring 2024, there were at least two other observations of adult *O. cornuta* on Vancouver Island. In March 2024, a taxonomist from the NBSBC observed a female and male *O. cornuta* about 15 km northwest of Black Creek on an artificial mason bee house alongside *O. lignaria* (Fig. 1B). In April 2024, a female *O. cornuta* was spotted in a mason bee nesting block in Comox, 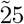 km from Black Creek. Together, these observations indicate multigenerational establishment of *O. cornuta* in British Columbia over a range of at least 170 km.

We tested whether visual features of cocoons, such as size and color, permitted the discrimination of *O. cornuta* from *O. lignaria* and *O. cornifrons. Osmia* cocoons have a broadly similar appearance, though there was apparent variability in size and color both within and across species (Fig. 3A). We used a standardized imaging pipeline to extract area, length, width, and average red, green, and blue color value for each cocoon (Fig. 3B). Cocoon areas overlapped across the three species (Fig. 3C), with ranges of 0.25-0.76 cm^2^ for *O. cornifrons* (n = 81), 0.30-0.92 cm^2^ for *O. lignaria* (n = 130), and 0.39-1.17 cm^2^ for *O. cornuta* (n = 106). *O. cornuta* female cocoons (mean: 0.89 +/- 0.15 cm^2^) were distinctly larger than all other groups, whereas the size of *O. cornuta* male cocoons (mean: 0.64 +/- 0.16 cm^2^) overlapped with *O. lignaria* females (mean: 0.70 +/- 0.13 cm^2^) (Fig. 3C). Cocoon aspect ratios (length/width) also overlapped across species (ranges: *O. cornuta:* 1.52-2.00, *O. lignaria*: 1.55-2.54, and *O. cornifrons*: 1.45-2.63.) The ranges of red, blue and green values for average cocoon color similarly overlapped (Fig. 3D). To test whether species could be classified by combining information across the data dimensions, we trained a support vector machine (SVM) on the extracted visual features. We evaluated the model using precision-recall curves for two classes, *O. lignaria* and *O. cornuta*, to determine whether the native *O. lignaria* could be separated from the non-natives, and whether *O. cornuta* could be reliably detected (Fig. 3E). An ideal classifier would achieve perfect (1.0) recall for *O. cornuta*, so that no instances of the species are missed. The highest precision of our best SVM model at 1.0 recall was 0.79. At 0.9 recall, *O. cornuta* precision was 0.87. The model performed more poorly for *O. lignaria*, which has features intermediate to the other two species tested (Fig. 3C-D). The highest precision for *O. lignaria* at a recall above 0.5 was 0.92 (at recall = 0.54), while at 0.9 recall, precision was 0.82. The sawtooth shape of the *O. lignaria* precision-recall curve (Fig. 3E) indicates sensitivity to changes in cross-validation dataset splits and possible overfitting to the relatively small set of images. The SVM marginally outperformed human sorters, as 79% of *O. lignaria* and *O. cornifrons* cocoons submitted by Oregon beekeepers were labeled accurately. These results demonstrate that sorting cocoons to species on the basis of visual features poses challenges, even when considering multiple features.

**Figure 3:**
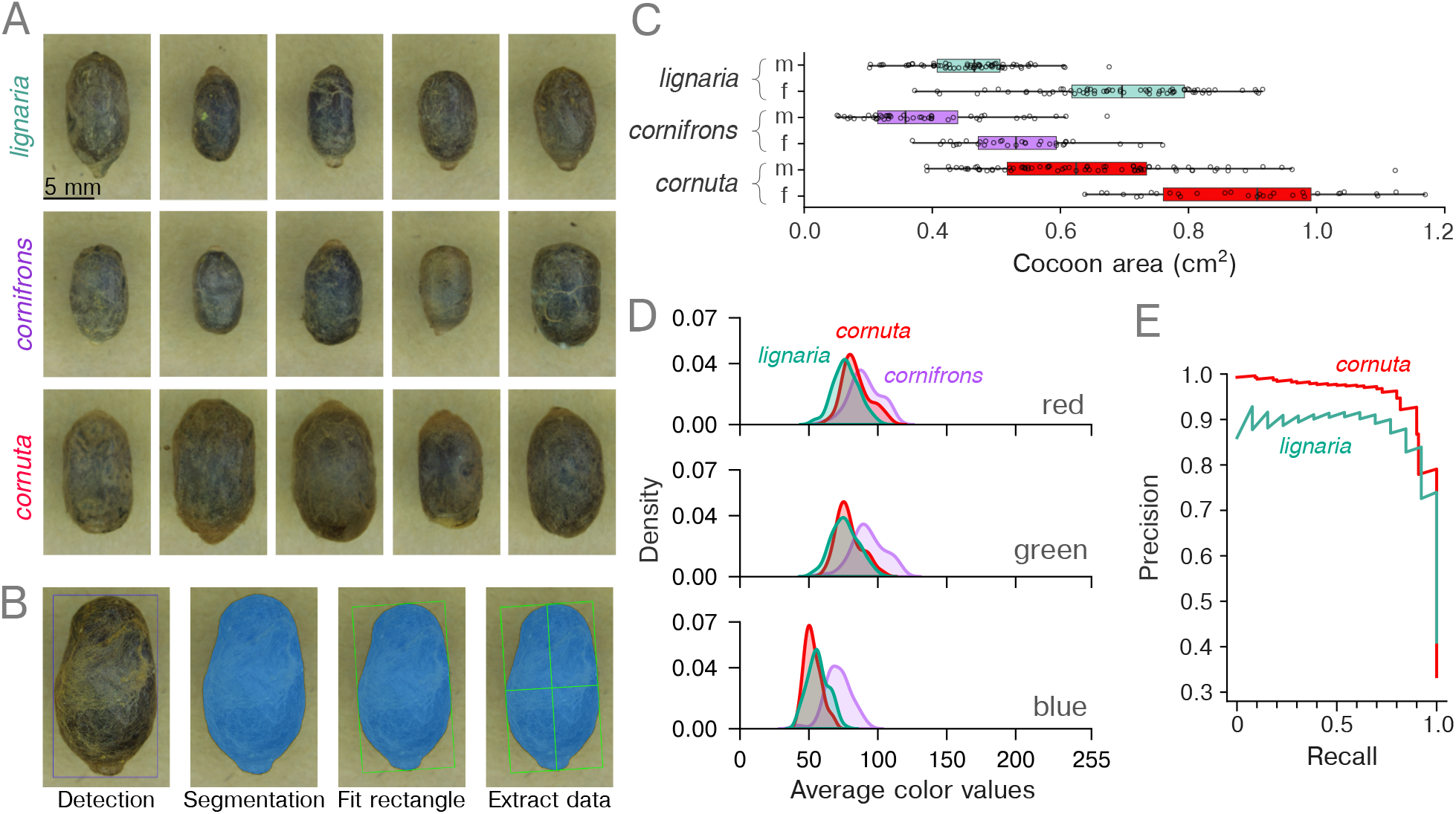
Visual features of *Osmia* cocoons provide a limited basis for interspecies discrimination. A) Randomly selected cocoon images for native *O. lignaria* (top row), *O. cornifrons* (middle row) and *O. cornuta* (bottom row). B) Pipeline for visual feature extraction involving automated detection, segmentation, and then fitting to extract length, width, area, and average color values. C) Measured cocoon areas for all three species, separated by sex. Box and whisker plot follows as in Fig. 2G. D) Smoothed histogram (via kernel density estimation) shows the distribution of average color values for each species measured on red, green, and blue channels. E) Averaged precision-recall curves for *O. lignaria* and *O. cornuta* obtained by training a support vector machine (SVM) model on the cocoon feature data over 1,000 cross-validated iterations. Precision is the ratio of true positives to total positive determinations. Recall is the proportion of actual positives correctly classified.

To evaluate potential range expansion dynamics for *O. cornuta*, we examined the historical pattern of North American establishment of the closely related non-natives *O. cornifrons* and *O. taurus*. Prior studies have shown that *O. cornifrons* and *O. taurus* widely expanded throughout the eastern US following introduction and establishment (Gutierrez et al. 2023, LeCroy et al. 2020, MacIvor et al. 2022).

We investigated the spatial pattern of *O. cornifrons* and *O. taurus* observations at a continent-wide scale, by mapping GBIF observations from 1960-2023 to a set of 918 equal-area bins (22,500 km^2^; 23-60 degrees latitude; Fig. 4A-C). During this period, observations of *O. lignaria* occurred across broad sections of North America, including the Pacific Northwest and southern British Columbia, the southwestern US, and much of the central and eastern U.S. (Fig. 4A). Observations of *O. cornifrons* were concentrated in the mid-Atlantic and northeastern U.S.; discontinuous records are found across the Great Plains, in Rocky Mountain states, and in all three Pacific coast states (Fig. 4B). Observations of *O. taurus* were similarly concentrated in the mid-Atlantic but there were isolated observations as far south as Jupiter Island, FL and west to Boise, ID (Fig. 4C). Since the advent of iNaturalist in 2008, there has been a marked increase in the total number and spatial range of occurrences for each species (Fig. 4D-F). Native *O. lignaria* and non-native *O. cornifrons* have been observed at similar annual rates, with a notably higher number of records for *O. taurus* (Fig. 4D). *O. lignaria* has been annually observed across a wider spatial range than *O. cornifrons* and *O. taurus* but this difference has narrowed in the last 5-10 years (Fig. 4E). Each of the three species continues to be observed in new locations for the first time, with close correspondence in rates of expansion between *O. cornifrons* and *O. taurus* (Fig. 4F). These data indicate *O. cornifrons* and *O. taurus* have been observed with increasing frequency over a broad, discontinuous spatial range since their respective introductions.

**Figure 4:**
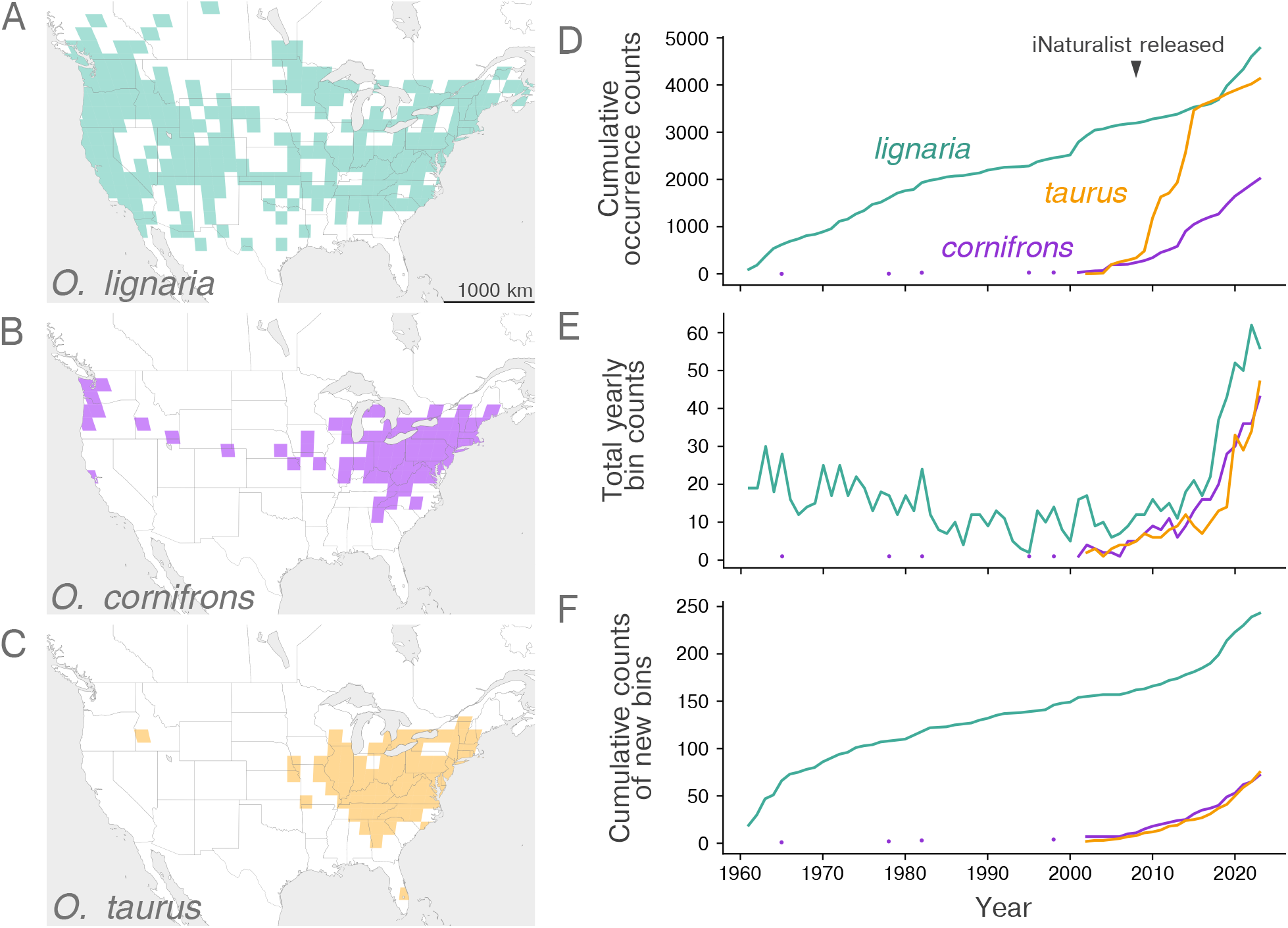
GBIF occurrence records suggest the spread of *O. cornifrons* and *O. taurus* in North America has been rapid yet discontinuous. A) Locations with GBIF occurrence records of the native *O. lignaria* from 1960 to 2023. Data are binned into equal-area regions (22,500 km^2^). B) Locations of bins with *O. cornifrons* records (data from 1965-2023). C) Locations of bins with *O. taurus* records (2002-2023). D) The cumulative count of confirmed GBIF observations for *O. lignaria, O. taurus*, and *O. cornifrons* from 1960-2023. Isolated records of *O. cornuta* between 1965-1998 are shown as points. E) The annual number of total bins in which each species was observed. F) The cumulative count of newly occupied bins (i.e. where a species was recorded for the first time).

Whereas O. *cornifrons* and *O. taurus* spread from the Mid-Atlantic, *O. cornuta* has established in the Pacific Northwest. To determine regions with suitable habitat for *O. cornuta* range expansion, we estimated the bee’s climate niche using occurrence data from its native range. We reduced 20 environmental variables (the 19 WorldClim bioclimatic variables and elevation) to four orthogonal principal components, which explained 85.2% of all native range variance (Fig. 5A, B). In the native range, the distribution of Mahalanobis distances (*D*^2^) to the centroid, was heavily right-skewed, ranging from 0.015 to 106.62 (unitless) with a mean of 4.0 (Fig. 5C, top axis). Based on this distribution, we set a suitability threshold at 61.65, which includes >99.9% of all values and removed 3 outliers. We calculated *D*^2^ to the native-range centroid across North America (24 - 55 degrees latitude; ∼1km^2^ resolution), to determine regions of potential suitability. North American *D*^2^ values ranged from 2.7 to 741.0 with a mean of 83.4 +/- 46.9 (Fig. 5C, bottom axis). 35.8% of locations had *D*^2^ values below the 61.65 suitability cutoff. There are two prominent regions in North America with *D*^2^ values below the native range threshold, indicating climate similarity (Fig. 5D). One region extends from Vancouver and Vancouver Island – the locations of *O. cornuta* detection – south to the Olympic Peninsula in Washington and the Willamette Valley in Oregon, inland through eastern Washington and northern Idaho through the Columbia River Valley, and through the major river valleys of southern British Columbia. A second, much larger region of high similarity in the central and eastern US ranges from the Ozarks in northwest Arkansas, through the Mid-Atlantic and north to Michigan, southern Ontario, New England, and Nova Scotia. The Mid-Atlantic, where *O. taurus* and *O. cornifrons* first established, exhibits some of the lowest *D*^2^ values in the range. The climate similarity map shows patchy, discontinuous regions of suitability throughout the American West, including the California Bay Area and in Utah near the Great Salt Lake. The site of one prior failed *O. cornuta* introduction in Logan, Utah is outside the range of estimated suitability, whereas another site, Dixon, California, is on the edge of estimated suitability. This model suggests that the Vancouver area was well suited to permit *O. cornuta* establishment, and that there is sufficient habitat for *O. cornuta* to spread throughout the Pacific Northwest and eastern North America.

**Figure 5:**
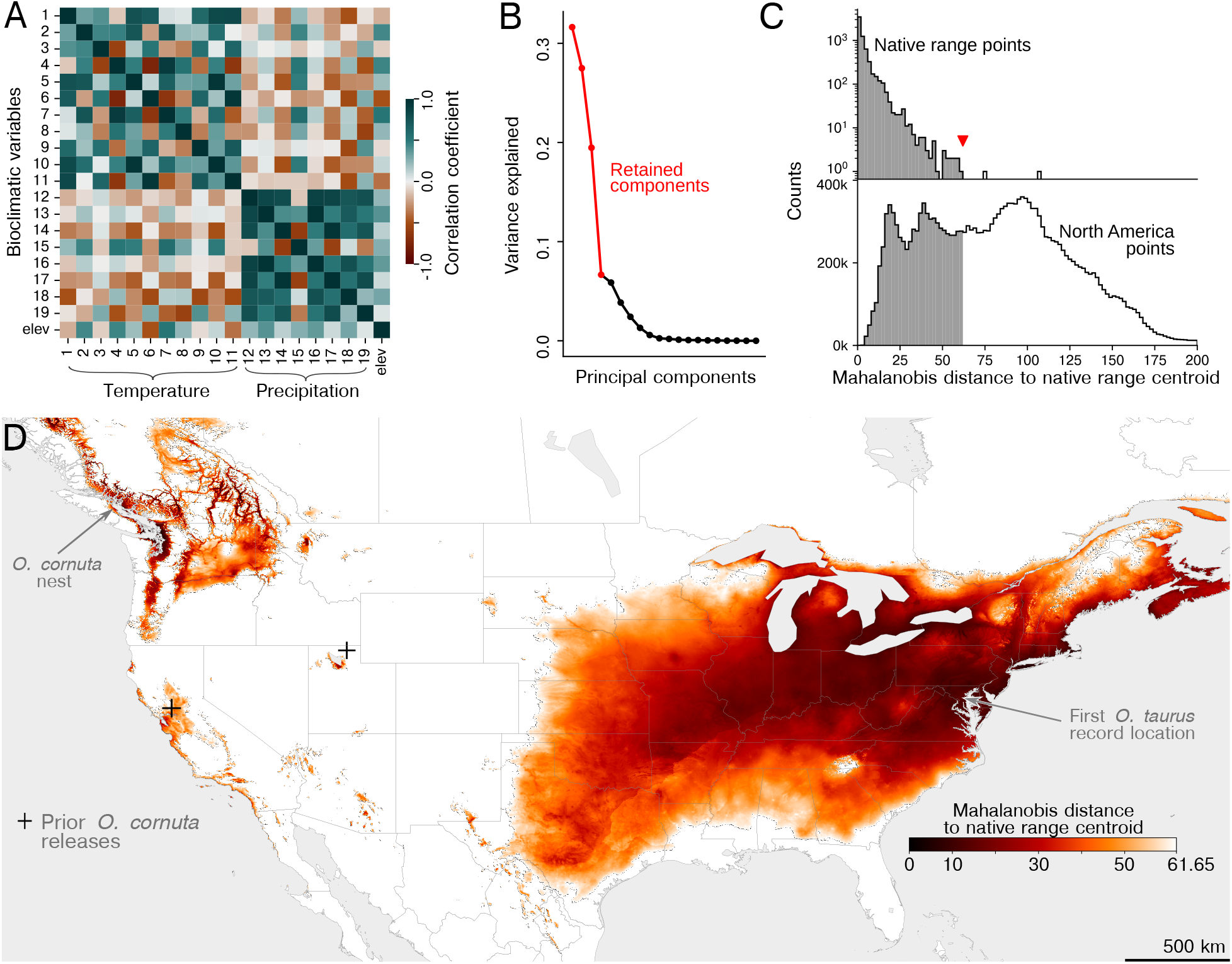
Large regions of North America have habitat similar to the native range of *O. cornuta*. A) Covariance matrix of data extracted at native range locations for the 19 WorldClim bioclimatic variables and elevation. B) Scree plot indicating variance explained by the 20 principal components calculated from the native range climate data. The first four components (in red) were used for subsequent analyses. C) Histograms of Mahalanobis distance (*D*^2^) distributions for the native range (top axis) and North American points (bottom axis). For both distributions, *D*^2^ was calculated to the native range centroid. The vertical axis for native range data is log-scaled to visualize the right tail of the distribution. The red arrowhead, (*D*^2^ = 61.65), denotes a habitat suitability threshold that encompasses >99.9% of native range points and excludes 3 outliers. D) Mahalanobis distance suitability model for *O. cornuta* in North America based on the WorldClim bioclimatic variables and elevation. Darker colors, with *D*^2^ closer to zero, indicate locations more similar to the native range. The Utah location where *O. cornuta* was previously released but did not establish is outside the range of predicted similarity. The first records of *O. cornuta* and *O. taurus* are in regions of high climate similarity.

## Discussion

In this study we documented multiple recent records in British Columbia, Canada of the European orchard bee, *Osmia cornuta*, a species previously thought to be limited to Europe and its periphery. These records, ranging over two years and 170 km, included a multigenerational nest containing hundreds of cocoons (Fig. 1). The physical features of these cocoons overlapped with those of other *Osmia* species (Fig. 2, 3), providing limited basis for classification (Fig. 3). This raises the possibility that *O. cornuta* will be inadvertently mixed into cocoon shipments of other mason bees. The rapid and discontinuous continentwide expansion of *O. cornifrons* and *O. taurus* further suggests the potential for anthropogenic dispersal of *O. cornuta* (Fig. 4). We estimated the climatic niche of *O. cornuta* using occurrence data in its native range and found that broad portions of the Pacific Northwest, southeastern Canada, and the central and eastern U.S. provide suitable habitat (Fig. 5). Given access to artificial dispersal vectors and favorable habitat, *O. cornuta* could spread quickly in North America.

The proximity of recent records to international ports suggests *O. cornuta* arrived in western Canada on commercial container ships. Across a wide variety of taxa, the introduction and invasion of foreign organisms has become more frequent with globalized trade (Bertelsmeier 2021, Fenn-Moltu et al. 2023, Perrings et al. 2005). This is especially the case with insects, which are small, mobile, and can stay dormant during development (Bertelsmeier 2021, Fenn-Moltu et al. 2023). Cavity-nesting *Osmia* s.s. are particularly prone to long-distance dispersal through shipping materials, where they can remain undetected as cocoons during transit (Russo 2016). The initial 2023 sighting of *O. cornuta* was only about 7 km from the Port of Vancouver, the fourth largest in North America. Subsequent Vancouver Island sightings were within 50-80 km of several international ports (e.g. Nanaimo, Alberni; Fig. 1B). Similarly, *O. taurus* was first detected in North America less than 30 km from the Port of Baltimore (Droege and Maffei 2023; Fig. 3B). The emergence of *O. cornuta* on Vancouver Island parallels the recent, well-publicized invasion by the the northern giant hornet (*Vespa mandarinia*), which was first encountered less than 5 km from the Port of Nanaimo (Zhu et al. 2020). This pattern of exotic insect detection near ports highlights the vulnerability of international commercial exchange points to invasion.

Our findings suggest that inadvertent shipments of cocoons could contribute to the future spread of *O. cornuta* in North America. The initial records of *O. cornuta* on Vancouver Island – in which adults and cocoons were observed alongside native *O. lignaria* – indicate the two species are likely to intermingle in shared nesting blocks. Once mixed, *Osmia* s.s. cocoons will be difficult to distinguish reliably on the basis of individual visual features such as size and color (Fig. 3). A support vector machine, which combines information across multiple features, performed better for *O. cornuta* than *O. lignaria*. Classifying *O. lignaria* with high precision is required to reliably sort non-native cocoons from native. The best performance the SVM obtained for *O. lignaria* was 0.91 precision at 0.54 recall (i.e. 1 in 10 identifications were wrong, and nearly half of the true positives were undetected). This exceeds the performance of volunteer beekeepers in Oregon; of 211 cocoons we received with *O. lignaria* or *O. cornifrons* labels, 79% were correctly identified. However, given that commercial *Osmia* rearing operations produce millions of cocoons a year, this performance would result in a significant number of errors and wasted cocoons, making it infeasible to control inadvertent dispersal of non-native species. Identifying *O. cornuta* for removal from mixed groups of cocoons would require perfect recall. For *O. cornuta*, the SVM achieved 1.0 recall, but only at a precision of 0.79. With this performance, one in five *O. lignaria* or *O. cornifrons* cocoons would be misclassified as *O. cornuta* and removed from the population. Alternative classification approaches (e.g. with a larger dataset or with a deep learning model) could potentially exhibit improved performance.

The rapid yet discontinuous range expansion of *O. taurus* and *O. cornifrons* in North America further supports the idea that *O. cornuta* could spread widely via anthropogenic dispersal. We found that the observed range of occurrences of non-natives *O. taurus* and *O. cornifrons* has continuously expanded in the past 20 years – with a notable range increase relative to a recent study (Gutierrez et al. 2023). For both species, the spatial pattern of observations was patchy, with isolated observations of *O. taurus* in Boise, ID and South Florida, and disparate records of *O. cornifrons* in Colorado, Utah, Idaho, and California (Fig. 4B-C). These isolated records may have resulted from long-distance cocoon shipments, and may serve as further case studies for possible establishment after translocation.

Our habitat niche model indicates broad regions of the Pacific Northwest and eastern North America have environmental conditions similar to *O. cornuta*’s native range. This similarity-based approach to identifying suitable habitat is not exhaustive; *O. cornuta* could succeed in climate conditions distinct from those in its native range or quickly adapt to new habitats. The analysis, however, highlights the most probable regions for *O. cornuta* spread, and provides a potential explanation for the failure of prior attempts to introduce *O. cornuta* to North America. The Logan, UT introduction site is outside the area of predicted suitable habitat, whereas the Dixon, CA site is just on the edge, indicating marginal suitability. The regions of predicted *O. cornuta* habitat suitability overlap conspicuously with the current ranges of closely related *O. lignaria, O. taurus*, and *O. cornifrons* (compare Fig. 4a-c, Fig. 5). If introduced to central or eastern North America (e.g. through the mason bee cocoon trade), it is possible *O. cornuta* will exhibit similar rapid expansion dynamics as its non-native congeners.

*O. cornuta* could affect native North American bees through pathogen transmission and direct competition for resources. *O. cornuta* are known to host *Chaetodactylus osmiae* mites which infest mason bee nest cells, *Monodontomerus obscurus* wasps which parasitize cocoons, and chalkbrood fungi (Krunic et al. 2005). *O. cornuta* are pollen generalists, with preferences for rosaceous trees and shrubs, *Salix, Quercus*, and *Acer* (Kratschmer et al. 2020) – several groups that are known preferences of *O. lignaria* (Kraemer and Favi 2005, Kraemer et al. 2014, Pinilla-Gallego and Isaacs 2018, Rust 1990). When coreleased in orchards, the pollen preferences of the two species were identical (Torchio and Asensio 1985). The similarities between *O. lignaria* and *O. cornuta* in size (Fig. 2b), biological requirements (Torchio and Asensio 1985, Bosch and Kemp 2003, Bosch and Kemp 2004), and diet suggest the two species could compete for nesting and floral resources where their ranges overlap.

Our study highlights the utility of community science monitoring, both for detecting biological invasions and predicting exotic species’ spread. Notably, most North American *O. cornuta* records were contributed by affiliates of the Native Bee Society of British Columbia; the cocoon collection was reported to the same organization. In many cases, community naturalist programs (e.g. Bee Atlases, Native Bee/Plant Societies, Natural Heritage Programs) have been shown to improve the quality of public records by increasing awareness of target species and sharing expertise through identification workshops, surveying events, or by moderating non-expert accessions (Kosmala et al. 2016, Anderson et al. 2020, Pocock et al. 2015). There are known limitations to grassroots reporting, such as spatial bias around human population centers (Amano and Sutherland 2013, Bowler et al. 2022) and potential taxonomic errors. The unsystematic nature of community science monitoring can also obscure historical trends in species distributions. Biodiversity monitoring capabilities have become more accessible and widespread in past decades, which is useful for monitoring the spread of a recently introduced species such as *O. taurus* or *O. cornifrons* (Fig. 4D-F). However, the increase in sightings of long-established natives such as *O. lignaria* is difficult to separate from the overall increase in sampling frequency and range. To estimate population size, assess range accurately, or create historical distribution benchmarks, targeted studies with standardized monitoring are essential.

Currently, *O. cornuta* is not listed as an invasive species of concern by Canada or the US. Policymakers should respond promptly to limit the spread of *O. cornuta* while implementing systematic monitoring efforts to determine the bee’s spatial range and ecological impact. Ideally, systematic surveys could operate in tandem with dispersed, public data collection. Community reporting will be a critical resource for tracking *O. cornuta* adults, which are easily identifiable by their bright red coloring and detectable by hobbyists with backyard mason bee houses. The commercial *Osmia* cocoon trade demands particular attention – both as a potential vector for unintended long-distance dispersal and also as a possible means for conducting large-scale monitoring. Effective, automated approaches for tracking exotic *Osmia* s.s. could generalize to aid in detecting other rare or invasive insects.

## Acknowledgements

We thank Seth Arendell for imaging, dissecting, and identifying the cocoons, Julia Jones, Lauren Ponisio, and Posy Busby for feedback on a draft version of this paper, and Weng-Keen Wong for discussions regarding analytical methods.

